# Pharmacodynamics of linezolid-plus-fosfomycin against vancomycin-susceptible and -resistant enterococci in vitro and in vivo of a Galleria mellonella larval infection model

**DOI:** 10.1101/624577

**Authors:** Caifen Qi, Shuangli Xu, Maomao Wu, Shuo Zhu, Yanyan Liu, Hong Huang, Guijun Zhang, Jiabin Li, Xiaohui Huang

**Author notes:** These authors contributed equally to this work (Caifen Qi, Shuangli Xu, Maomao Wu). Correspondence: Xiaohui Huang, Department of Basic and Clinical Pharmacology, School of Pharmacy, Anhui, Medical University, Meishan Road, 81,230032, Hefei, Anhui, China, Jiabin Li, Department of Infectious Diseases, The First Affiliated Hospital of Anhui Medical, University, Jixi Road, 218, Hefei, Anhui, China.

## Abstract

**Objective:** To explore the in vitro and in vivo antibacterial activity of linezolid/fosfomycin combination against vancomycin-susceptible and -resistant enterococci (VSE and VRE), providing theoretical basis for the treatment of VRE.

**Methods:** The checkerboard method and time-kill curve study were used to evaluate the synergistic effect of linezolid combined with fosfomycin against VSE and VRE. The transmission electron microscopy (TEM) was employed to observe the bacterial cell morphology followed by each drug alone and in combination, elucidating the possible result of antibiotic combination therapy. The Galleria mellonella infection model was constructed to demonstrate the in vivo efficacy of linezolid plus fosfomycin for VSE and VRE infection.

**Results:** The fractional inhibitory concentration index (FICI) values of all strains suggested that linezolid showed synergy or additivity in combination with fosfomycin against five of the six strains. Time-kill experiments demonstrated that the combination of linezolid-fosfomycin at 1×MIC or 2×MIC led to higher degree of bacterial killing without regrowth for all isolates tested than each monotherapy. TEM imaging showed that the combination treatment damaged the bacterial cell morphology more obviously than each drug alone. In the Galleria mellonella infection model, the enhanced survival rate of the combination treatment was revealed compared to linezolid monotherapy (P<0.05).

**Conclusions:** Our data manifest that the combination of linezolid and fosfomycin may be a possible therapeutic regimen for VRE infection. The combination displays excellent bacterial killing and inhibits amplification of fosfomycin-resistant subpopulations.

## 1. Introduction

Once considered a part of the normal gastrointestinal flora, enterococcus species have emerged as the second leading cause of healthcare-acquired infections in the United States, now associated with life-threatening infections such as pyelonephritis, intra-abdominal infections and bloodstream infections (1). They are intrinsically resistant to most commonly used antibiotics and readily acquire resistance (2). The VRE clinical isolate was first reported in 1988 in New York, from a wound culture (3). Resistance to vancomycin is primarily mediated via acquisition of transferrable plasmids encoding modification of the primary binding site D-Ala-D-Ala. These peptidoglycan precursors are replaced with D-Ala-D-lactate or D-Ala-D-Serine and vancomycin loses its affinity by approximately 1000-fold (4). VRE has been associated with 2.5 times higher mortality compared with VSE, which might be associated with postponed appropriate antibiotic therapy (5, 6). Infections caused by VRE, incidences of which have been on the rise since 1988, poses significant challenges for infection treatment because there are fewer and fewer available antimicrobial agents (7). VRE has become problematic in the clinical setting due to the given the tendency for easy spreading and challenges in antimicrobial management (8).

Linezolid, which was approved by US Food and Drug Administration in 2000, is recommended as one of the first-line antimicrobial agents for the treatment of VRE infection (9). Linezolid is bacteriostatic, and adverse events attributed to the long-time use of linezolid such as neurotoxicity and bone marrow toxicity limit its use (10, 11). In addition, Smith et al reported that prolonged exposure of linezolid increased the likelihood of the emergence of linezolid-resistant Enterococcus faecium (12). Fosfomycin is a bactericide against both Gram-positive and Gram-negative bacteria, including E. faecium (13). Fosfomycin has currently aroused interest as a potential therapeutic choice for infections caused by VRE despite limited efficacy data (14, 15). The rapid occurrence of fosfomycin resistance in vitro is the dominating limiting factor for its use as monotherapy in the clinical practice (16), which results in this old antibiotic often being considered specifically for use in combination with another agent.

In this study, we described the in vitro antimicrobial activity of linezolid in combination with fosfomycin against clinical VSE and VRE. We also evaluated the efficacy of this regimen in vivo using the Galleria mellonella infection model. Our findings highlighted the potential of this combination for treating infections caused by VRE.

## 2. Materials and Methods

### 2.1 Bacterial Isolates

Six strains were studied, including type strain vancomycin-susceptible (ATCC 29212) and -resistant (ATCC 51299) E. faecalis and vancomycin-susceptible (No.1) and -resistant (No.2, No.3, No.4) E. faecium. ATCC 29212, ATCC51299 and No.1 were supplied by the First Affiliated Hospital of Anhui Medical University. No.2, No.3, No.4 were obtained from Beijing Hospital.

### 2.2 Antimicrobial Agents and Medium

Linezolid was obtained from Pfizer limited liability company (Shanghai, China). Vancomycin and fosfomycin were purchased from National Institutes for Food and Drug Control. Antibiotic stock solutions were freshly prepared in Milli-Q water (Labconco Corporation), which were sterilized by a 0.22-um sterilizing filter (MET, the United States) each day (fosfomycin), or were reserved at −20 °C and used within a month (linezolid 1280 μg/mL, vancomycin 1280 μg/mL).

Cation-adjusted Mueller–Hinton broth (CAMHB, Oxoid, England), containing calcium of 25mg/L and magnesium of 12.5 mg/L was used for all in vitro susceptibility analyses. Mueller-Hinton agar (MHA, Oxoid, England) was used for culturing bacteria, agar dilution method and quantifying colony counts.

### 2.3 Determination of Antimicrobial Susceptibility Testing

Minimum inhibitory concentrations (MICs) of all antibiotics except fosfomycin were determined using broth microdilution methods. Bacteria which were cultured to the log-phase (approximately 1.5×10^8^ CFU/mL) and then diluted 150-fold, were seeded at 96-well plates which were added a series of two-fold dilutions of antimicrobial agents. Plates were incubated in humidified 5% CO_2_ at 37 °C for 24h. After the incubation period, the lowest concentration of antibiotics at which no visible bacteria grew was determined as MIC. MIC of fosfomycin was detected by agar dilution method, using MHA containing a series of two-fold diluted fosfomycin appended with glucose-6-phosphate of 25 μg/mL. The results were interpreted in the light of the MIC breakpoints of the Clinical and Laboratory Standards Institute (CLSI, 2018) antimicrobial susceptibility testing standards (17). ATCC29212 was served as a quality control strain. All experiments were repeated three times.

### 2.4 Checkerboard Assays

The chequerboard broth microdilution assay was performed in 96-well microtitre plates with 2-fold dilutions of two antibiotics which were diluted in CAMHB. Linezolid ranging from 1/64×MIC to 2×MIC was dispensed in every row, whereafter, fosfomycin supplemented with 25 μg/mL of glucose-6-phosphate ranging between 1/64×MIC and 2×MIC was added in each column. An equal volume of standardized bacterial suspension of 1×10^6^ CFU/mL was added and then all plates were incubated at 37°C in an aerobic atmosphere for 24h. Fractional inhibitory concentrations (FICs) were calculated as the MIC of drug A or B in combination divided by the MIC of drug A or B alone, respectively, and the FIC index (FICI) was obtained by adding the two FIC values. To categorise the drug combination that consistently generated the lowest FICI after repeating the experiment in duplicate on two further occasions, the results can be grouped as follows: FICIs of ≤ 0.5 were interpreted as synergistic; FICIs of >0.5 but ≤1 were considered as additive; FICIs of >1 but <4 were considered as no interaction; and FICIs ≥4 were interpreted as antagonistic. SBPI was also calculated as previously described (18). Combinations demonstrated to be synergistic or additive were assessed using the time-kill methods.

### 2.5 Time-Kill Studies

Time-kill studies were performed on ATCC2912, No.1, No.2, No.4 isolates based on a previously reported method (19). The concentration of linezolid and fosfomycin was selected on the basis of drug serum concentration in stable state that is achievable when administrated the optimal dosage. In brief, bacterial suspensions at the exponential-phase were diluted to the inoculum of approximately 5.0×10^5^ CFU/mL in 8 mL of fresh CAMHB in 15 mL of tubes with variable concentrations of linezolid (0.5×, 1× and 2×MIC) and fosfomycin (0.5×, 1× and 2× MIC) alone or in combination. The tubes were then incubated with shaking at 37 °C. At 0, 2, 4, 6, 8, 12 and 24 hours, the bacteria in each tube were diluted with 4 °C 0.9% sodium chloride, and then seeded on MHA plates for viable colony counts. Synergy, additive effect, indifference and antagonism were defined as ≥2 log_10_ CFU/mL kill, <2 but >1 log_10_ CFU/mL kill, ±1 log_10_ CFU/mL kill, and >1 log_10_ CFU/mL growth, respectively (19). Bactericidal activity was defined as 99.9% reduction in cell numbers from the initial inoculum. Changes to fosfomycin MICs were measured for all strains that regrew after 24h to detect whether these strains were resistant to fosfomycin.

### 2.6 Transmission Electron Microscopy (TEM)

We employed TEM to explore the influence of the linezolid-plus-fosfomycin on the cellular structure and morphology of vancomycin-susceptible enterococcus faecium No.1 and Vancomycin-resistant enterococcus faecium No.2. Bacteria which were cultured to the logarithmic phase were transferred and then diluted 100-fold into the tube which contained 2 mg/L linezolid, 128 mg/L fosfomycin, or both antibiotics, continuing culturing for 4h in CAMHB in the light of the time-kill experiments. Samples were centrifuged for 10 min at 3300 rpm and 4 °C three times. Supernatants were discarded and bacteria in the bottom of the tube washed with 1mL phosphate-buffered saline (PBS) during centrifugation procedures. After the final centrifugation procedure, the supernatants were abandoned, and then bacterial pellets resuspended and fixed in 1 mL PBS with 2.5% glutaraldehyde at 4℃ overnight. After fixed, tubes were centrifuged at 3300 rpm for 10 min, the fixed agent removed, and bacterial pellets washed three times in 1 mL PBS as above, and then dehydrated gradiently with 30%, 50%, 70%, 80%, 90% and 100% ethanol. Each time, it was placed for 15 minutes and centrifuged for 10 min at 3300rpm. At last, bacterial pellets were washed with 100% ethanol twice as above, and then resuspended in 1mL 100% ethanol. The prepared samples were observed under TEM at Southeast University, China.

### 2.7 Galleria mellonella Infection Model

The Galleria mellonella infection model was constructed according to a previously reported method with slight variations (20). G. mellonella larvae were stored in the darkness at 2-10℃ and were used within 7 days of receipt. Larvae weighing 250mg-350mg, milky white and active, without grey marks were selected for all experiments. The bacterial suspension at log-growth phase was centrifuged, washed and resuspended in 0.9% NaCl three times. All inocula were determined by bacterial colony counts on MHA. In order to determine 80% lethal doses of No.1 and No.2, eight G. mellonella larvae of each group were injected with 10 μl bacterial suspension of three different concentrations of 10-fold dilution using a 25 μl Hamilton microliter syringe via the last left proleg. Larvae in Petri dishes were reared at 37℃ in an aerobic and humid atmosphere and were observed every 24h until 96h. Larvae whose body was blackening and showed no movement in response to touch was considered dead. The doses of linezolid and fosfomycin were calculated according to the doses administered in the human body. 96 larvae were randomly selected and equally assigned to each of the following six groups: (i) linezolid alone (10 mg/kg), (ii) fosfomycin alone (200 mg/kg), (iii) linezolid (10 mg/kg) and fosfomycin (200 mg/kg) in combination, (iv) linezolid (5 mg/kg) and fosfomycin (100 mg/kg) in combination; (v) linezolid (2.5 mg/kg) and fosfomycin (50 mg/kg) in combination; or (vi) no treatment. Larvae were inoculated with 80% lethal doses of either No.1 or No.2 as above, followed by 10 μl injections of the test drug or 0.9% NaCl as a control within 2h after injection, and then observed as performed above. Treatment was given only once. Blank and 0.9% NaCl controls were set for each experiment. The results of any experiment with more than one dead larva in either control group were abandoned. All experiments were performed twice on different occasions.

### 2.8 Statistical Analysis

Survival curves were constructed using the Kaplan–Meier method and compared by log-rank test using GraphPad Prism 5. The charts in the figures were plotted using GraphPad Prism 5. For all experiments, P-values <0.05 were considered statistically significant.

## 3. Results

### 3.1 Antimicrobial Susceptibility Testing

The results of in vitro susceptibility testing are listed in Table 1.The MICs of linezolid against all six tested strains ranged from 1 to 4 μg/mL. MICs of all organisms to fosfomycin were 128 μg/mL. In short, no bacteria were resistant to linezolid and fosfomycin.

**Table 1.** MICs of vancomycin, linezolid, fosfomycin and linezolid-fosfomycin combination against six enterococci strains

| | <sup>a</sup>MIC(μg/mL) | | | | | |
| --- | --- | --- | --- | --- | --- | --- |
|  | VAN | LIN | FOS | LIN+FOS | FICI | SBPI |
| ATCC29212 | 2 | 2 | 128 | 1+64 | 1.0 | 3 |
| ATCC51299 | 256 | 2 | 128 | 0.03125+64 | 0.52 | 65 |
| No.1 | 1 | 2 | 128 | 0.5+32 | 0.5 | 6 |
| No.2 | 512 | 2 | 128 | 0.5+32 | 0.5 | 6 |
| No.3 | 256 | 1 | 128 | 0.25+128 | 1.125 | 8.5 |
| No.4 | 512 | 4 | 128 | 1+64 | 0.75 | 3 |

MIC, minimum inhibitory concentration; VAN, vancomycin; LIN, linezolid; FOS, fosfomycin;
FICI, fractional inhibitory concentration index; SBPI: susceptible breakpoint index
a, Antibiotic susceptibility breakpoints:
VAN: $\leq 8$ mg/mL, susceptible (S); 8-16 mg/mL, intermediate (I); $\geq 32$ mg/mL, resistant (R).
LIN: $\leq 2$ mg/mL, susceptible (S); 4 mg/mL, intermediate (I); $\geq 8$ mg/mL, resistant (R).
FOS: $\leq 64$ mg/mL, susceptible (S); 128 mg/mL, intermediate (I); $\geq 256$ mg/mL, resistant (R).

### 3.2 In Vitro Synergy Testing with the Chequerboard Method

The fractional inhibitory concentration index (FICI) values of all strains suggested that linezolid showed synergy or additivity in combination with fosfomycin against five of the six strains (Table 1). No antagonistic effect was observed against all isolates evaluated. For No.1 and No.2, the existence of fosfomycin at 0.25×MIC reduced the MIC of linezolid from 2 μg/mL to 0.5 μg/mL. A FICI of ≤0.5 was seen for both strains, demonstrating a synergistic interaction. No significant synergism was observed in ATCC29212, ATCC51299, No.3 and No.4. However, a SBPI >2 was discovered in all six isolates tested, which manifested potential synergism (Table 1).

### 3.3 Time-Kill Studies

The results of time-kill studies are shown in Figure 1. Linezolid at 1×MIC showed bacteriostatic activity to all four strains. For VSE ATCC29212, No.1, fosfomycin at 1×MIC resulted in 2.1, 2.4 log_10_ CFU/mL colony decrease at 8h or 12h, respectively; Another, fosfomycin at 1×MIC generated 0.8, 1.6 log_10_ CFU/mL reduction in bacterial growth after 8 hours incubation against No.2, No.4, respectively. However, regrowth appeared in all four isolates after 24h and paralleled the growth of the controls in two strains. Fosfomycin resistance was noted at 24h when used as monotherapy. MICs for all isolates were >1024 mg/L, representing at least a eight-fold MIC elevation.

**Figure 1.**
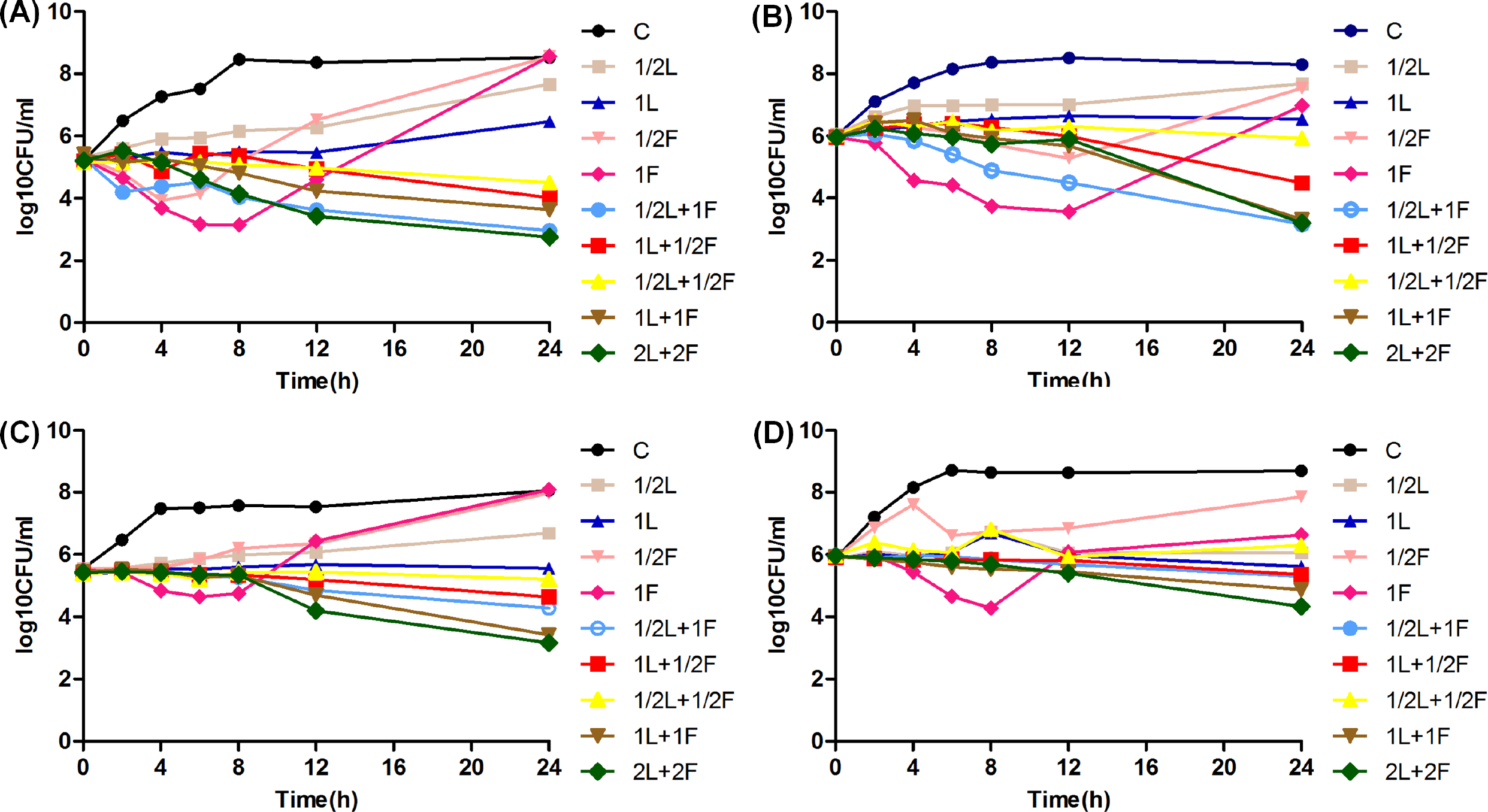
time-kill study performed on vancomycin-susceptible enterococcus faecalis type strain (A, ATCC 29212), vancomycin-susceptible enterococcus faecium (B, No.1), vancomycin-resistant enterococcus faecium (C, No.2. D, No.4), L, linezolid; F, fosfomycin; C, control group.

On the contrary, linezolid in combination with fosfomycin showed better bacterial killing activity without regrowth for each of the isolates in comparison with any agent alone. The combination treatment at 1×MIC demonstrated synergistic bacterial killing against No.1 and No.2, and also produced an additive effect against ATCC29212 and No.4. The combination of 2×MIC was considerably more active than the combination of 1×MIC for all four strains evaluated.

### 3.4 Influence of Linezolid and Fosfomycin Alone and In Combination on the Cell Morphology of Vancomycin-Susceptible and -Resistant Enterococcus Faecium

Figure 2 showed TEM results of vancomycin-susceptible enterococcus faecium No.1 and vancomycin-resistant enterococcus faecium No.2 following therapy with linezolid (1×MIC), fosfomycin (1×MIC), or both. For No.1, without treatment, the cells were observed with elliptical shapes with integrated cell membranes (Figure 2A). Both linezolid monotherapy and fosfomycin monotherapy resulted in significantly longer cell compared to the untreated group and appeared to be undergoing cell division (Figure 2B, 2C). In the combination treatment group, the bacterial cell membrane surface became disruptive (Figure 2D). For No.2, the cell morphology of the untreated cells was normal with round shapes with unbroken cell membranes (Figure 2E). Linezolid monotherapy had minimal impact on the gross morphology of the bacterial cells compared with the control group (Figure 2F). Compared to the untreated group, the bacterial cell treated with fosfomycin monotherapy displayed uneven and rough shape with cell length increased to approximately double (Figure 2G). Linezolid/fosfomycin combination led to obvious cell membrane damage with leakage of cell cytoplasm (Figure 2H).

**Figure 2.**
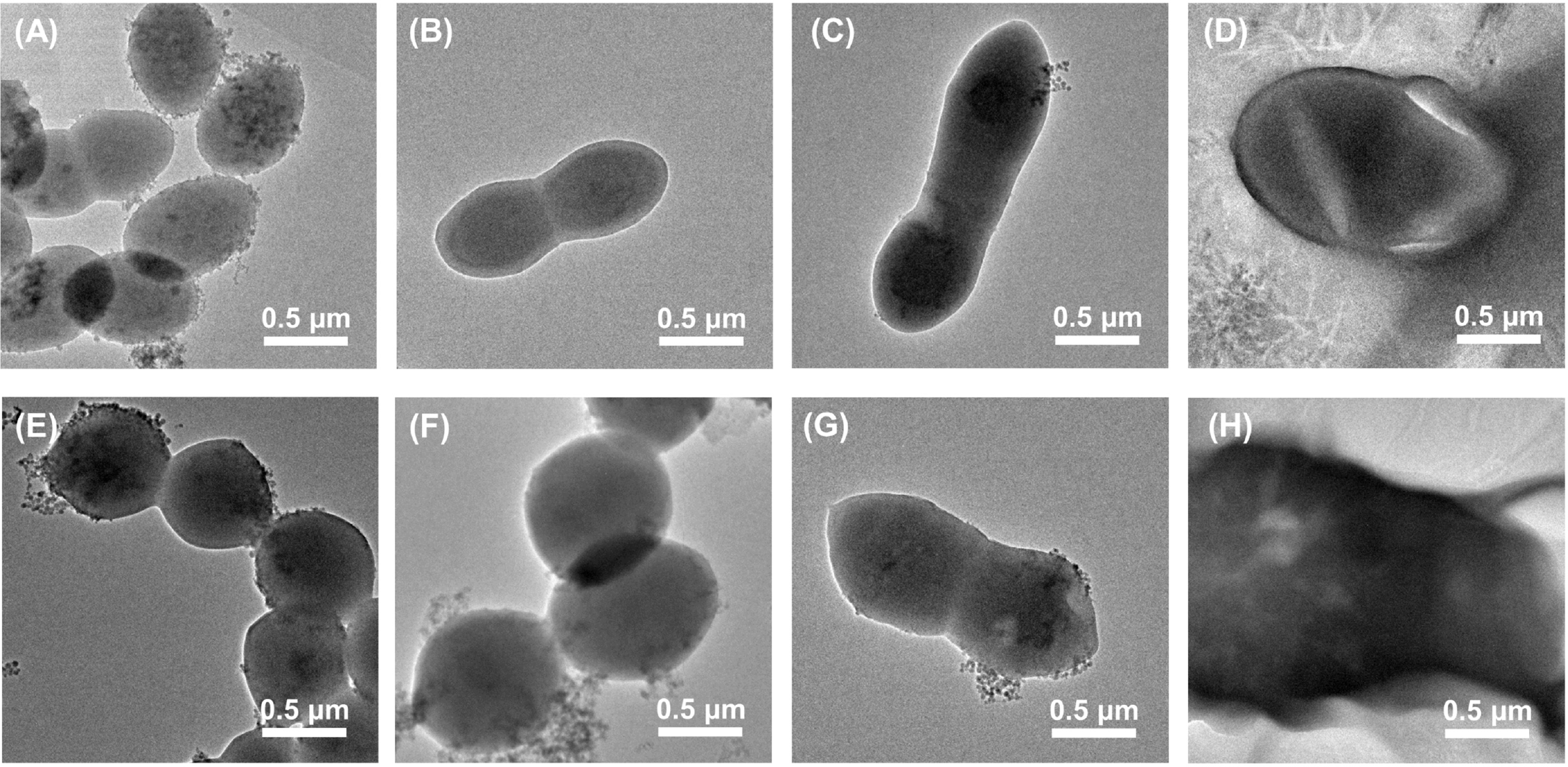
images from transmission electron microscopy for No.1 (B, C, D) and No.2 (F, G, H) treated with 2 mg/L linezolid, 128 mg/L fosfomycin, or both. A and E represent the control group.

### 3.5 Activities of Linezolid and Fosfomycin in Infected Galleria mellonella Larvae

As the bacterial concentration increased, the mortality of the Galleria mellonella larvae also increased, and most of the deaths of the infected larvae occurred within the first 24 hours. The 80% lethal dose of No.1 and No.2 was approximately 2×10^7^ CFU/mL and 10^8^ CFU/mL, respectively (Figure 3).

**Figure 3.**
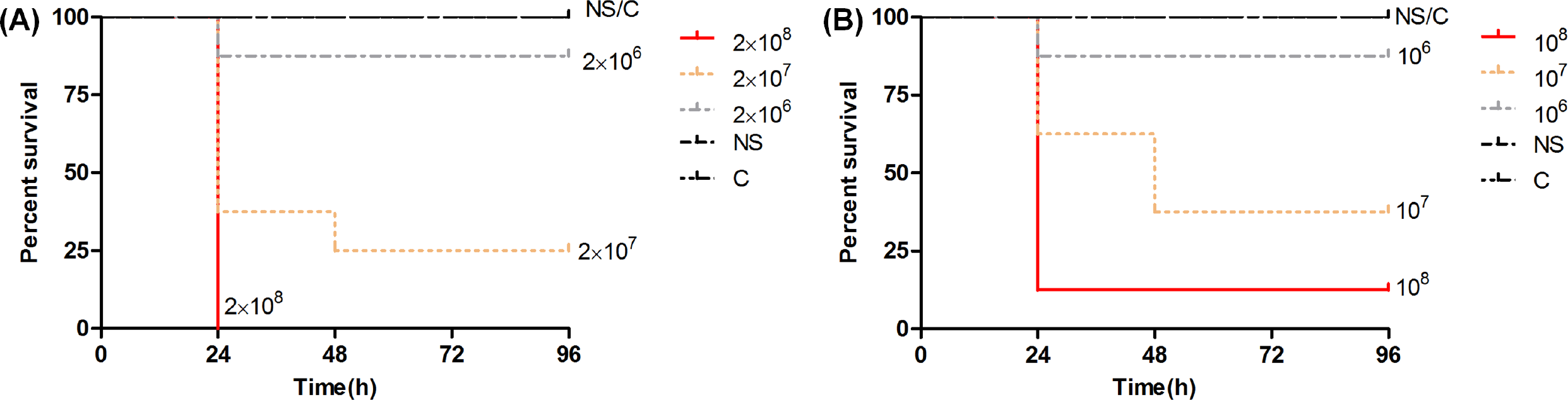
80% lethal dose of No.1 and No.2 in infected Galleria mellonella larvae. NS, 0.9% sodium chloride; C, control group.

The survival rate of linezolid against No.1 and No.2 was 62.5% and 43.75%, respectively. The combination of high doses was superior to the combination of low doses but no significance was observed. A statistically significant higher survival rate was observed for No.1 and No.2 in the combination of linezolid and fosfomycin compared with linezolid monotherapy (P <0.05). Interestingly, fosfomycin showed excellent antibacterial efficacy against both No.1 and No.2, which was approximately equivalent to the linezolid plus fosfomycin (P >0.05) (Figure 4).

**Figure 4.**
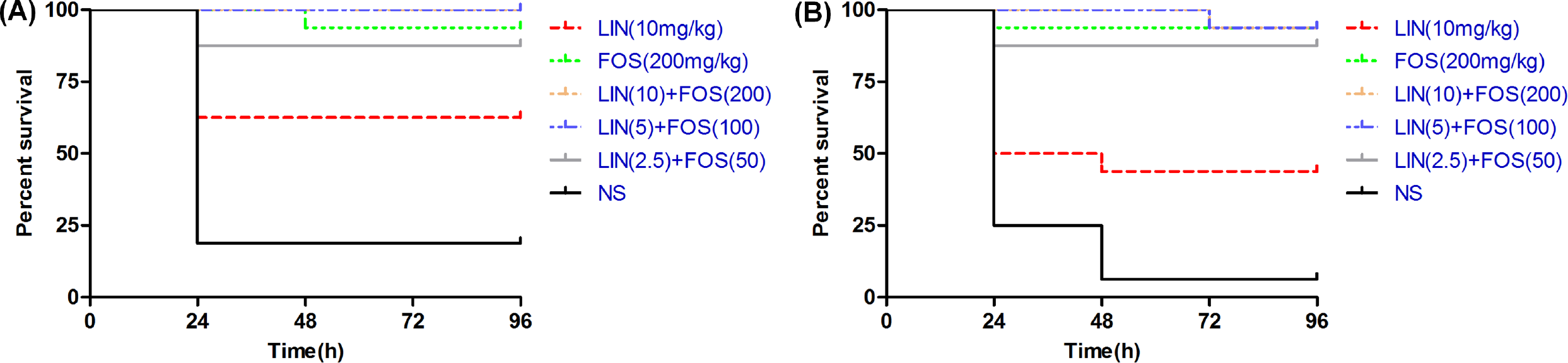
efficacy of linezolid alone, fosfomycin alone, and the combination of different doses against No.1 and No.2 in a Galleria mellonella infection model. LIN, linezolid; FOS, fosfomycin; NS, 0.9% sodium chloride.

## Discussion

Infections caused by VRE are on the rise in recent years worldwide, complicating the therapeutic options (1, 21). The increasing use of linezolid, one of the last-resort antibiotics, enhances the selective pressure for developing resistance to it in VRE strains (22). Currently, the speed of research and development of novel antimicrobial agents cannot keep pace with the increasing antibiotic resistance rates, so more and more unconventional combinations for the infection of VRE seem to be attractive options (23). The study conducted by Luther et al showed that the combination of linezolid and gentamicin combination enhanced antimicrobial activity against VRE (20). Tang et al reported that teicoplanin combined with fosfomycin revealed excellent synergistic activity against VRE (24). Previous studies have confirmed the potent synergism of the combination of linezolid and fosfomycin against another common multidrug-resistant gram-positive pathogen, methicillin-resistant staphylococcus aureus (MRSA) (25). However, reports in regard to this combination therapy when used against VRE are scarce.

In the current study, the FICI values of all strains suggested that linezolid showed synergy or additivity in combination with fosfomycin against five of the six strains (Table 1). No antagonistic effect was observed against all isolates evaluated. However, a SBPI >2 was discovered in all strains. SBPI is a parameter predicting the effect of antimicrobial combination therapies. Additionally, due to in-depth analysis of the pharmacokinetic/pharmacodynamic indicator, SBPI is likely to more related to clinical outcome compared to FICI. A SBPI >2 indicates synergy (18). In the time-kill curve, linezolid only displayed bacteriostatic liveness, in agreement with the results demonstrated by Oliva et al (26). For this reason, linezolid monotherapy is associated with higher failure rates for severe VRE infections, especially VRE bloodstream infections (27). Fosfomycin initially exhibited excellent bacterial killing, followed by regrowth after 8h or 12h, which is consistent with that reported in a recent study conducted in an experimental foreign-body infection model (26). This phenomenon may be interpreted by “fosfomycin heteroresistance”. Heteroresistance to fosfomycin has been exhibited in Streptococcus pneumoniae and MurA (UDP-*N*-acetylglucosamine enolpyruvyl transferase) where is responsible for the heteroresistance (28). There is a growing body of evidence suggesting that resistance to fosfomycin can emerge with monotherapy (16, 26). The high-level resistance of enterococcus to fosfomycin may result from mutations of the target enzyme MurA, accompanied by a slight decrease in catalytic activity (29). Therefore, fosfomycin monotherapy is problematic for treating infections caused by VRE. In contrast, the combination of linezolid with fosfomycin at 1×MIC or 2×MIC resulted in a higher degree of bacterial kill without regrowth for each of the isolates than either monotherapy regimen. The combination treatment at 1×MIC showed a synergistic effect against No.1 and No.2, and also displayed an additive activity against ATCC29212 and No.4. The synergism of the linezolid plus fosfomycin was described in several studies, especially against MRSA (25, 30). Based on my research, only one study was found that explored the effectiveness of linezolid combined with fosfomycin against VRE in vitro via time-kill curve experimentation and similar results were detected in that study (31). Fosfomycin inactivates MurA via covalently combining to the thiol group of a cysteine located in the active site of MurA, causing the early synthesis of the peptidoglycan precursor of bacterial cell wall to be suppressed and therefor causing bacterial death (15). According to the above study, it is speculated that the mechanism for the augmented bacterial killing revealed in the combination is inhibition of bacterial cell wall biosynthesis by fosfomycin, which results in easier entry of linezolid into bacterial cells.

TEM imaging results showed the damage of the bacteria was more obvious in the combination treatment group, which further confirmed the synergistic effect of linezolid combined with fosfomycin and tentatively clarified the mechanism of action of the combination. After retrieving similar investigations, it appears our study was the first to observe the impact of linezolid combined with fosfomycin on VRE’s cell morphology using TEM, and the results are in line with the above-mentioned results of in vitro.

The Galleria mellonella larva infection model has been previously used for the research of the virulence of numerous human pathogens and the efficacy of the antimicrobial agents (32–34). G. mellonella has also been effectively employed to test the effects of rifampicin combination therapy against enterococcal infections in the past(35). When compared with mammalian models, G. mellonella is cheaper to obtain and free of ethical constraints (36). Additionally, G. mellonella possesses both cellular and humoral immune response, which functions analogously to vertebrate immune systems (37). In the study, the combination of linezolid and fosfomycin improved survival rate significantly over linezolid alone, while no significant difference was detected between the combination treatment group and fosfomycin group. This result was in accord with the results in vitro and might preliminarily predict clinical outcomes and indicated that combination therapies with linezolid plus fosfomycin might be a good therapeutic option for serious VRE infections.

In conclusion, linezolid combined with fosfomycin has good in vitro and in vivo activity against VSE and VRE in contrast to linezolid or fosfomycin monotherapy. Importantly, the combination also inhibits amplification of fosfomycin-resistant subpopulations. Even so, further mammal experiments and clinical studies are needed to confirm the activity of this combination on VRE and the exact mechanism of the combination.

## Acknowledgments

This study was supported by the National Natural Science Fund of China (0601021203); the Fund of Excellent Talents in Colleges and Universities of Anhui Province, China (gxbjZD06); and the Fund of Academic Leaders of Anhui Province, China (2015D068).

